# In an arms race between host and parasite, a lungworm’s ability to infect a toad is determined by host susceptibility, not parasite preference

**DOI:** 10.1101/2021.10.19.464902

**Authors:** Harrison JF. Eyck, Gregory P. Brown, Lee A. Rollins, Richard Shine

## Abstract

Evolutionary arms races can alter both parasite infectivity and host resistance, and it is difficult to separate the effects of these twin determinants of infection outcomes. Using a co-introduced, invasive host-parasite system (the lungworm *Rhabdias pseudosphaerocephala* and the cane toad *Rhinella marina),* we quantified behavioural responses of parasite larvae to skin-chemical cues of toads from different invasive populations, and rates at which hosts became infected following standardised exposure to lungworms. Chemical cues from toad skin altered host-seeking behaviour by parasites, similarly among populations. The number of infection attempts (parasite larvae entering the host’s body) also did not differ between populations, but rates of successful infection (establishment of adult worm in host lungs) was higher for range-edge toads than for range-core conspecifics. Thus, lower resistance to parasite infection in range-edge toads appears to be due to less effective immune defences of the host rather than differential behavioural responses of the parasite. In this ongoing host-parasite arms-race, changing outcomes appear to be driven by shifts in host immunocompetence.

## 1. Introduction

Biological invasions offer ideal systems with which to study host-parasite co-evolution (1, 2). Colonisation of novel habitat destabilises co-evolutionary relationships, allowing us to study reciprocal antagonistic selection and its consequences (3, 4). As invasive hosts expand into novel habitat, processes such as genetic drift, diversifying selection (5) and the consequences of dispersal (e.g., spatial sorting) (6), drive the emergence of phenotypic differences among subpopulations. In turn, such divergences can rapidly generate variation in a parasite’s traits, including infectivity (7), causing geographic and temporal mosaics in host-parasite traits (2). In this way, invasions can substantially alter the dynamics of co-evolutionary relationships (8, 9). However, teasing apart these temporally and spatially dynamic relationships remains challenging.

An arms race can impose strong selection on traits that affect infectivity and resistance (10), potentially modifying the interaction between host and parasite at multiple phases in their interaction. Parasites with a direct life cycle (i.e. without an intermediate host) must initially locate a host. To this end, many parasites show specific (11, 12) and flexible behaviours (13–15), that optimise host-seeking. Once located, a parasite must penetrate the host’s peripheral defences, including its physical outer barrier and microbiome (16–18), and circumvent avoidance behaviours by the host (19). Finally, parasites must withstand the host’s immune defences. Hosts may invest heavily into functions such as antimicrobial capacity of blood plasma (20), whereas parasites evolve to cloak themselves against such defences (21, 22). In total, changes in infection rates during an arms-race might evolve via a range of mechanisms that include traits of the parasite (e.g., host-seeking), and/or the host (e.g., peripheral barriers, immune defences).

One intensively studied biological invasion is that of the cane toad *(Rhinella marina)* in tropical Australia. Native to South America, toads were introduced to Hawai’i to control sugar cane beetles. In 1935, 101 individuals were transported from Hawai’i and bred in Queensland, where their offspring were released along a 1200km stretch of the Queensland coastline (23). The toads’ spread across Australia has proven unstoppable (24). Evolutionary processes such as natural selection and spatial sorting have generated morphological, physiological, and behavioural differences between range-edge (Western Australia; WA) and range-core populations (Queensland; QLD) (Reviewed in; 25). A native-range parasite, the nematode lungworm *Rhabdias pseudosphaerocephala* (hereafter lungworms), survived the toad’s multiple translocations (26) and occurs across the Australian range of the toad. An arms-race has developed between toads and lungworms (27), resulting in toads from different parts of the invasive range having differential resistance to infection (30). Potential explanations for that spatial variation in vulnerability include differential skin permeability (28), attractiveness to lungworm larvae (29), or investment in immune responses (30) in toads from different areas. This system presents a robust opportunity to tease apart adaptations and the parasite *versus* its host in driving divergence in infectivity during an arms-race.

In this study, we investigate whether lowered resistance to lungworms in toads from range-edge populations within the Australian invasive range is due to shifts at three phases of infection: 1) host-attraction, 2) peripheral defences, or 3) immune defences. First, we exposed geographically intermediate Northern Territory (NT) lungworm larvae to chemical cues from the skins of toads from three different populations (WA, NT, and QLD) to quantify the effect of those cues on activity levels of lungworm larvae. Then we exposed toads from all three populations to infective NT lungworm larvae, recording the number of lungworms that attempted infection, and the number of lungworms that successfully penetrated the lungs. If differences in infection resistance are due to differences in attraction, then lungworm activity rates should be higher in the presence of cues from range-core toads. In contrast, if peripheral defences of the host are the main barrier to infection, we expect lungworms to attempt infection more often on WA toads. Finally, if immune defences of the host drive differences in infection resistance, the proportion of lungworms that succeed in penetrating the lungs (relative to infection attempts) should be higher in range-edge hosts than in toads from other populations.

## 2. Materials and methods

### (a) Lungworms

Rhabditid lungworms have a heterogonic life cycle. Hermaphroditic adults reside in the lungs, where they deposit eggs that are carried by mucus to the mouth and swallowed into the digestive system. These eggs hatch into sexually reproductive L1 adults which exit the host in the faeces, where they develop and reproduce. The offspring of the L1 adults, known as L3, hatch within the L1 mother and feed on her organs for 4 to 7 days, after which they burst out of her body. L3 larvae are infectious; they seek out hosts, burrow through their tissue to reach the lungs and develop into adults.

### (b) Generation of toads and lungworms

For infection trials, we extracted eggs from 50 adult lungworms taken from a single, wild adult toad from the Northern Territory. We placed these eggs into a petri dish in a water droplet using a pipette. We extracted faeces from the colon of this same toad to provide food for larvae.

For the ethogram, we took eggs from three adult lungworms from a single, wild adult toad, captured in the Northern Territory (different from the toads used in infection trials). We placed eggs in individual petri dishes, such that all the eggs from each individual adult worm were grouped together. We took faeces from a non-infected Northern Territory toad to be used as food for each of the three populations of lungworm L1 larvae, to produce L3 for the experiment.

We used 30 toads in total for both experiments, comprising 10 metamorphs from each region, bred from adults that had been collected in Fitzroy Crossing in WA (range-edge; toads present < 10 years), Pine Creek in NT (intermediate range; toads present for ≈ 20 years), and Townsville in QLD (range-core; toads present for > 80 years: see (24). All metamorphs weighed approximately five grams each.

### (c) Host-searching behaviour of lungworm larvae

To quantify how parasites change their behaviour in response to cues from hosts, we exposed lungworms to toad cues on agar. To create agar plates, we began by mixing 2g of 2.0-4.5% agar ash (Sigma-Aldrich A7002) with 98ml of water and bringing this to boil for 1 min. While still liquid, this mixture was pipetted into sterile 35mm diameter petri dishes and left for 45-60 min to set. We then kept a toad on each plate overnight to impart chemical cues onto the agar. Toads were then removed, and to begin a trial, we added a 2.5μl water droplet containing a single lungworm to the plate at a point in the centre. We recorded the behaviours of lungworms using a microscope camera (MEE-500, Yegren Optics, China). We recorded for 10 minutes in each trial and analysed the videos using ImageJ v1.52p (31), measuring movement speed and time spent moving/still.

### (d) Infection trials

For the infection trials, we created fresh agar plates in the same manner as described above. We collected lungworms in 10μl of water using a pipette, placed ten onto each agar plate, and waited ten minutes for acclimation. Afterwards, we added one toad to the centre of each agar plate and began the trial. We exposed toads to lungworms six times each, with fresh agar plates each time. We conducted trials over the course of four days. In each time-period we also used ten control plates which we treated identically to the trials, but without toads present. We gave toads a minimum of 30 min rest between trials. Once trials concluded, we removed toads from the agar plate and placed them into their holding tanks. We flooded used plates with 1ml of water and swirled the water vigorously side to side, counting lungworms as they were removed from the plate with a pipette. At 15-16 days after trials were finished, we euthanised metamorphs in MS-222 (Tricaine methanesulfonate) and the adult lungworms were collected from toad lungs and counted.

### (e) Statistical analysis

We conducted all statistical analyses using R version 4.0.2 (32), with packages lme4 v1.1-26 (33) and MASS v7.3-54 (34). Normality and heteroskedasticity were assessed by visually observing residual plots, qq-plots, and density plots of model residuals. Non-normal models were either fit with a suitable distribution in a generalised linear model or log-transformed to satisfy model assumptions. Differences between groups were assessed using analysis of variance (ANOVA) from the package car v3.0-10 (35).

## 3. Results

### (a) Host-searching behaviour of lungworm larvae

#### Does the presence of toad cues affect activity of L3s?

Exposure to toad cues did not significantly influence the proportion of time lungworms spent moving *(χ^2^* = 1.89, *P* = 0.17), but reduced their average speed *(χ^2^* = 10.84, *P* < 0.01; Fig. 1).

**Figure 1.**
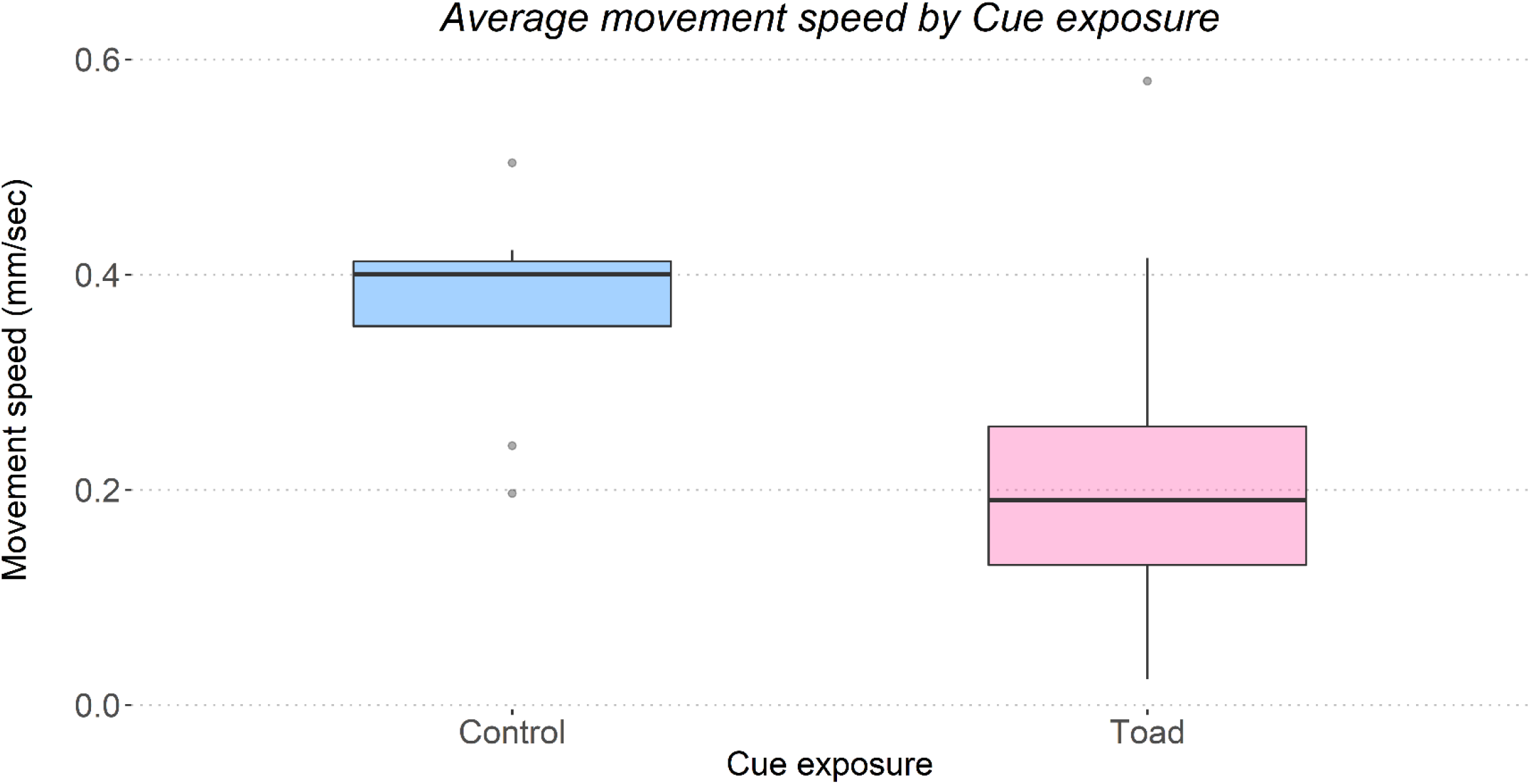
Box and whisker plot showing absence or presence of toad (Rhinella marina) cue on the x-axis and average movement speed of lungworm (Rhabdias pseudosphaerocephala) larvae in mm/second on the y-axis. Worm larvae exposed to toad cues moved more slowly (χ^2^ = 6.45, P = 0.04). N = 8 larvae exposed to controls, 25 larvae exposed to cues from toads.

**Figure 1.**
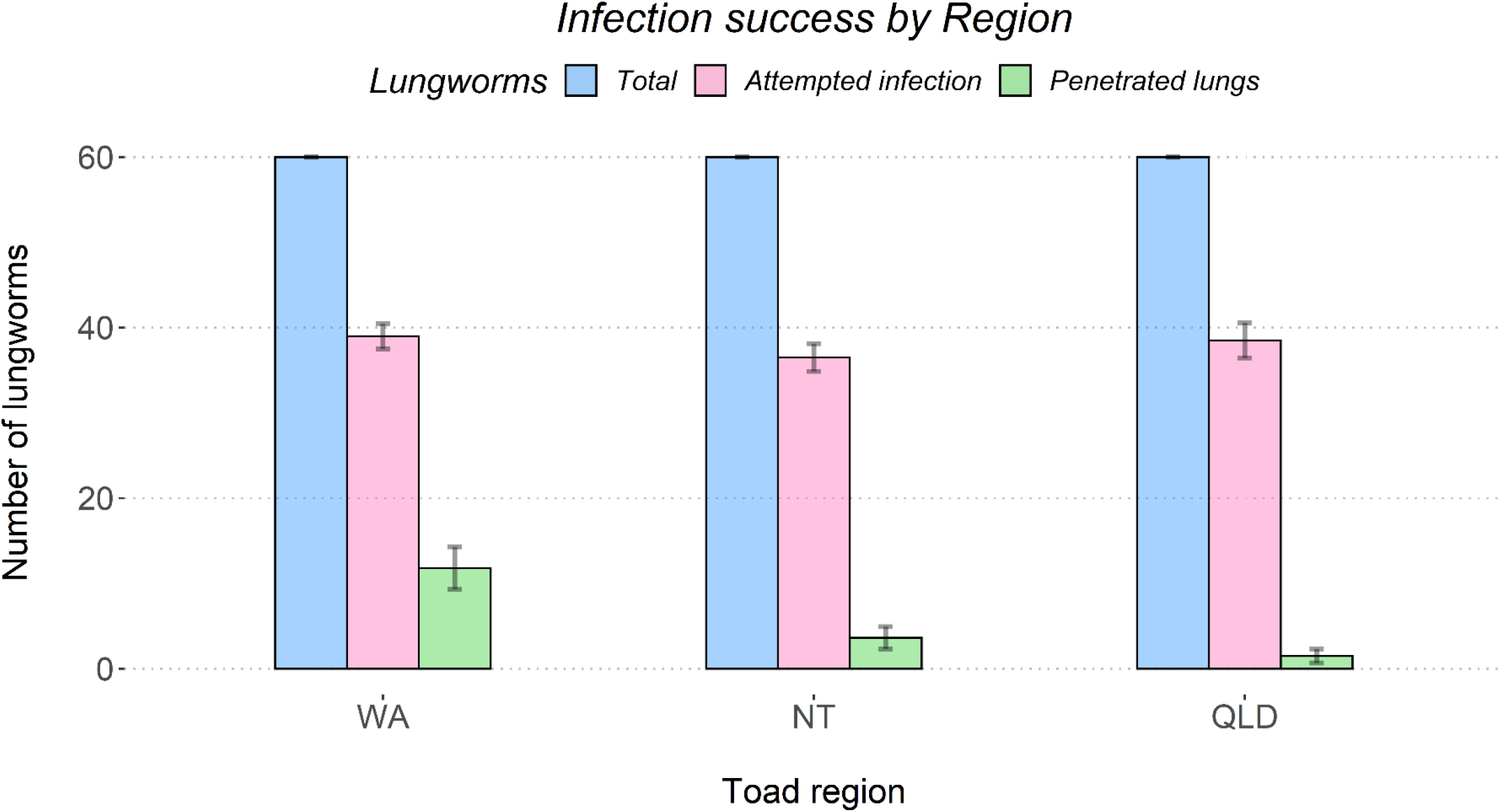
Bar plot showing the mean number of lungworms (Rhabdias pseudosphaerocephala) from the Northern Territory that toads (Rhinella marina); were exposed to in total (blue), that attempted infection (pink), and that successfully penetrated the lungs of metamorph cane toads (green). The plot is grouped by toad origin on the x-axis, with the count of lungworms shown on the y-axis. The plot shows infection attempts did not differ by region (pink bars, χ^2^ = 0.79, P = 0.67), but successful infection attempts did (green bars, χ^2^ = 14.47, P < 0.01), with WA toads being particularly susceptible to infection. QLD = Queensland; NT = Northern territory; WA = Western Australia.

#### Does attractiveness differ among toads from different regions?

The region from which a toad cue was sourced had no significant influence on the proportion of time lungworms spent moving *(χ^2^* = 0.73, *P* = 0.69) nor lungworm movement speed (*F_2,19_* = 2.95, *P* = 0.08).

### (b) Infection trials

#### Peripheral defences

Neither the average movement speed of lungworm larvae *(χ^2^* = 1.13, *P* = 0.29) nor the region from which toads were sourced *(χ^2^* = 1.67, *P* = 0.43; Fig. 2) significantly influenced the number of cumulative infection attempts made by lungworms.

#### Internal defences

Toad region influenced the success rate of lungworms infection establishment (i.e., penetrated the lungs: *χ^2^* = 14.47, P < 0.01. Fig. 2), with range-edge toads three to ten times more susceptible to infection than were conspecifics sourced from other areas. The average rates that lungworms successfully penetrated the lungs of toads, from all worms that attempted infection, were 30.26% in WA, 9.93% in NT, and 3.90% in QLD (Fig. 2).

## 4. Discussion

Our results clarify how a parasite has gained an advantage over its host during an evolutionary arms-race. Lungworms exhibited similar behavioural responses (activity levels and number of infection attempts) when they encountered range-edge and range-core toads, yet were far more successful in establishing infections in range-edge (WA) toads than in toads from intermediate or range-core areas. The behavioural response of lungworm larvae to chemical cues from the host – slowing down – likely enhances the parasite’s ability of encountering a mobile host and initiating attempts at penetration (36). Lungworm larvae in our study spent much of their time motionless, a common behaviour in parasitic nematodes that ambush their hosts (37–40), and moved more slowly than usual when exposed to host cues. Different cues may stimulate different behaviours (41–43). The environment where L3 larvae and toads most commonly interact is likely within toad burrows, because toads spend a lot of time inside them, reuse them (44), and L3 numbers inside burrows are increased by the accumulation of faeces, and the relatively cool moist conditions. By moving infrequently and slowing down when a toad is in the local area, lungworms may be better able to locate a host and infect it.

Importantly, the behavioural response of lungworm larvae to toad cues was not significantly affected by the geographic origin of the toads; and nor was the number of larvae that attempted to penetrate the toad’s body. These results suggest that the major phase of host-parasite interactions affected by the arms race is the ability of lungworm larvae to travel through the host’s body and establish infections in the lungs, rather than earlier phases such as recognising host presence through chemical cues or initiating attempts to penetrate the host’s body. Consistent with our results, previous research has shown that blood plasma from range-edge (WA) toads is relatively ineffective at killing range-core worms (7), and that lungworms from the range-edge are larger than are those from other locations (27). Such local adaptation and differences in lungworm infectivity between populations may explain why range-edge toads are resistant to some populations (7), yet were susceptible to the NT lungworms used in this study. Toads at the range-edge thus may be exposed to different kinds of lungworms, as well as fewer of them, favouring investment into different kinds of immune defences such as eosinophils, macrophages, or inflammation.

Changes in the evolved interaction between toads and their lungworm parasites appear to have been especially rapid at the toad invasion’s range-edge in WA (7, 29), likely because low population densities of hosts (toads) at the invasion front decrease rates of parasite transmission (27). As a result, selection for mounting an effective immune response to fight parasite attack may be weaker at the invasion front than elsewhere (consistent with the enemy release hypothesis: (45). The level of energy investment into immunocompetence also may trade off with energy allocation to dispersal, a trait under strong evolutionary pressure (both from natural selection and spatial sorting) at an expanding range edge (30). In combination, these pressures may generate a toad with phenotypic traits that render it more susceptible to infection if attacked by a parasite: there is little advantage to costly immune defences if resources are better invested in dispersal (46, 47). Future studies on *R. pseudosphaerocephala* could further investigate differences in infectivity amongst different lungworm populations. Additionally, future research could use a gene expression approach to identify genomic regions that respond to infection in toads and investigate how these differ across host regions.

## Ethics

This research was conducted under ethics approval from Macquarie University Animal Research Authority (ARA) (reference no. 2021/001) to GPB and RS.

## Data accessibility

The raw data and R code used for analysis are provided in electronic supplementary material.

## Authors’ contributions

The study was designed with input from all authors. HJFE collected the lungworms and ran the experiments, GPB provided the toads used in the experiments. HJFE analysed the data with comments from GPB, RS, and LR. HJFE and RS wrote the manuscript with comments from GPB and LR. All authors approved the final version and agree to be held accountable for the content.

## Acknowledgements

We thank our respective laboratories and institutions for making this research possible.

