## Supplementary methods for "In an arms race between host and parasite, a lungworm’s ability to infect a toad is determined by host susceptibility, not parasite preference"

1 Methods
1.1 Generation of lungworm larvae
1.2 Host-searching behaviour of lungworm larvae
1.3 Infection trials
2 Analysis
2.1 Host-searching behaviour of lungworm larvae
2.2 Infection trials
3 Supplementary table 1: model results and sample sizes
4 Supplementary material references

### 1. Methods

#### 1.1 Generation of lungworm larvae

For the infection trials, we removed 50 adult lungworms by dissection from the lungs of a single, wild adult cane toad caught from Marlow Lagoon, Northern Territory (-12.48° S, 130.96° E). We removed the lungworm eggs by poking a hole in the tissues of the adult lungworm using a sharp metal blade, this causes the eggs to burst out as the high-pressure inner cavity of the worm is exposed to the low-pressure environment. We placed these eggs into a petri dish in a water droplet using a pipette. We extracted faeces from the colon of the same toad the lungworms were taken from and placed it in the petri dish in water at room temperature with natural day/night cycles, to provide food for developing L1 larvae. L3 infective larvae emerge after approximately four days of cultivation.

For the behaviour trials, we took three adult lungworms from a single adult toad captured in the wild from Marlow Lagoon, Northern Territory (-12.48° S, 130.96° E). We removed eggs from these worms and placed them in individual petri dishes, such that all the eggs from each individual adult worm were in their own petri dish. We took faeces from a non-infected Northern Territory toad and thoroughly searched it for larvae under a dissecting microscope at 4x magnification. When no larvae were found, these faeces were used as food for each of the three populations of lungworm L1 larvae.

We used 30 toads in total for both experiments, comprising 10 metamorphs from each region, bred from adults that had been collected in Fitzroy Crossing in WA (range-edge; toads present < 10 years), Pine Creek in NT (intermediate range; toads present for ≈ 20 years), and Townsville in QLD (range-core; toads present for > 80 years: see (24). All metamorphs weighed approximately five grams each. These toads were given individual identification numbers and housed separately in one-litre plastic tubs with 50ml of water provided and changed daily.

#### 1.2 Host-searching behaviour of lungworm larvae

To quantify how lungworms change their behaviour in response to hosts, we exposed the larvae to toad cues on agar. To create agar plates, we began by mixing 2g of 2.0-4.5% agar ash (Sigma-Aldrich A7002) with 98ml of water and bringing this to boil for 1 minute. While still liquid, this mixture was pipetted into sterile 35mm diameter petri dishes and left for 45-60 minutes to set. To infuse the assay plate with the toad cue, live toads (the same individuals as used in the infection trials) were housed overnight for one night on the agar plates with water added. Toads were not fed for the duration of the trials, to remove the possibility they would defecate on infection plates, preserving a level of cue equality across plates. Metamorph toads were removed in the morning and the behaviour assays took place on that same day. This meant there were ten plates with WA toad cue, ten plates with QLD toad cue and ten plates with NT toad cue.

To begin the trial, we added a single worm to the plate at the dot in the centre in a 2.5µl droplet of water. Once the water evaporated and freed the worm, the trial began. We recorded the behaviours of the worms using a microscope camera (MEE-500, Yegren Optics, China). To ensure that worms stayed in-frame, an observer was present for every trial to move the plate as the worms moved. We recorded behaviour for 10 min in each trial. One of the worms on a control plates escaped underneath the agar and was not included in the data. As a result the sample size for the control trials was nine. We recorded behaviours using the video footage generated from filming using the program ImageJ v1.52p (1). Worm behaviour was classed into either movement or still. For movement we recorded time and distance, with speed calculated from these measurements. For each "still" phase, we recorded the amount of time spent immobile. We recorded the level of magnification for each trial to calibrate distance measurements. The distances of each behaviour were recorded using the segmented line tool in ImageJ, where the distance of the behaviour was the distance between the tip of the larvae's posterior end at the beginning of the movement, to the tip of the larvae's posterior end at the conclusion of the movement. Four of these individual lungworm trails ran short (ID numbers X2.2N, X1.3Q, X3.2Q, X2.2W), as the individual lungworms were lost briefly during filming. The time they were lost was excluded from the analysis, but data on average speed and proportion of time spent moving measured from the time these individuals were tracked was retained in the analysis.

#### 1.3 Infection trials

For the infection trials, we created fresh agar plates in the same manner as mentioned above, but no toad cue was deposited onto the agar. Agar plated were left in the refrigerator overnight, and the following day we collected ten L3 stage larvae in 10µl of water using a pipette and placed the larvae on the surface of each petri dish. We counted larvae as they were collected and counted once again as they were deposited on the agar to ensure accuracy. L3 larvae are capable of fast movement on agar plates such as these. Once every plate contained ten L3 larvae, we left them for ten minutes as an acclimation period. At this point, we added the toads to the centre of the agar plate, one toad per plate, and started the timer. We ran the trial six times per toad. We conducted trials over the course of four days. We used ten control plates in each of the six trials. We treated them identically to the trials apart from having no toads on the agar. Each trial was run separately, and we always gave toads a minimum of 30 min rest between trials. We recorded both the time of day the trial began and ambient temperature at the time of the trial. Once the time had expired, we removed the toads from the agar plate and placed them back into their holding tanks. To count how many L3 larvae remained on the plate after the assay, we flooded the plate with 1ml of water, and swirled the water vigorously side to side, clockwise and anti-clockwise. Larvae were counted twice each trial, once as we drew them up with a pipette and again as we ejected them into a separate, clean petri dish filled with water. L3 larvae have poor movement in water and are easy to count and collect in this manner.

After the trials had ended, we continued to provide water and food (wild-collected termites) to metamorphs daily. 15-16 days after trials were finished, metamorphs were euthanised in MS-222 (Tricaine methanesulfonate) and the now-adult lungworms were collected from their lungs and counted.

### 2. Statistical analysis

We conducted all statistical analyses using packages implemented in R version v4.0.2 (2). Models were constructed and assessed using the package lme4 v1.1-26 (3). Mixed models with grouping variables were used where appropriate. Where a singular fit occurred in a mixed model, a fixed effects model was used instead (4). Differences between groups were assessed using analysis of variance (ANOVA) from the package car v3.0-10 (5). Normality and heteroskedasticity were assessed by visually observing residual plots, qq-plots and density plots of model residuals. Non-normal models were fit with a suitable distribution in a generalised linear model to satisfy model assumptions. Sample sizes for each analysis and model results are shown in supplementary Table 1.

#### 2.1 Host-searching behaviour of lungworm larvae

*Does the presence of toad cues affect activity of L3s?*

We modelled the influence of toad cue on the proportion of time lungworms spent moving with a generalised linear fixed effects model using the glm function in the lme4 package. With proportion of time worms spent moving as the response variable, this model followed a quasibinomial distribution with treatment (toad cue vs control) and worm family ID grouping variable as fixed factors, worm length and ambient temperature as covariates.

We analysed whether or not toad cue influenced average movement speed of worms with a linear mixed effects model that had average lungworm movement speed as the response variable, treatment (toad cue vs control) fixed factor, worm length and ambient temperature as covariates, and worm family as a random intercept. If lungworms did not move at all in their trial, they were excluded from this analysis.

*Does attractiveness differ among toads from different regions?*

To determine if the proportion of time lungworms spent moving differed between toad regions, we created a generalised linear fixed effects model with the glm function in the lme4 package. This model followed a quasibinomial distribution with the proportion of time worms spent moving as the response variable. It had toad region of origin and the worm family ID grouping variable as fixed factors, and worm length and ambient temperature as covariates.

To establish whether average movement speed of lungworms was influenced by toad region-of-origin, we generated a linear fixed effects model with average movement speed as the response variable. Toad region-of-origin and worm family ID were fixed factors, with worm length and ambient temperature as covariates.

#### 2.2 Infection analysis

Supplementary material

To verify that our method of recovery from agar plates was accurate, we calculated the average number of lungworms successfully collected from control agar plates, where 10 lungworms were added to a plate and no toads were present for the duration of the trial. The success rate for these 59 trials in total was 97.80% ± 0.80.

To establish whether the number of worms recovered from agar plates differed between toad regions, or was affected by worm activity, we calculated the average speed of lungworms in the behavioural trial associated with each individual toad's cue and factored that into the infection attempt data for each metamorph. As the same toads were used to generate cue as were used in the infection trial, we used these data to determine if there was a relationship between lungworm activity and infection attempts for these toads. Worm recovery count from each individual toad cue plate was modelled by fitting a generalised linear mixed effects model using the glmer function in the lme4 package following a poisson distribution with a log link. Average speed of lungworms exposed to each individual toad cue, and region-of-origin of the toad were fixed effects, with individual toad ID included as a random intercept.

To determine if the number of worms successfully penetrating the lungs differed between populations of toad, the success rate of lungworms (successful infections/total number of infection attempts) was used as a response variable was fitted with a quasibinomial distribution in a generalised linear model, using the glm function from the lme4 package, with region-of-origin of the toad as the fixed factor.

Supplementary material

### 3. Supplementary table 1

*Supplementary table 1* showing sample sizes and model results for each model in the analysis. For the effect of toad region on successful infection attempts, sample size shows the total replicates, with total number of individual toads in parentheses. Significant effects highlighted in bold and italics. QLD = Queensland; NT = Northern territory; WA = Western Australia

| **Model** | ***Control n*** | ***Toad n*** | | **Predictor variable** | **χ2/F** | **Pr value** |
|---|---|---|---|---|---|---|
| *Proportion of time spent moving (Control v Toad)* | 9 | 30 | | Exposure (Toad v Control) | 1.89 | 0.17 |
| | | | | Worm length | 0.01 | 0.93 |
| | | | | Ambient temperature | 0.06 | 0.80 |
| | | | | Worm family | 0.21 | 0.90 |
| *Average movement speed (Control v Toad)* | 8 | 26 | | ***Exposure (Toad v Control)*** | ***10.84*** | ***< 0.01*** |
| | | | | Worm length | 0.10 | 0.76 |
| | | | | Ambient temperature | 0.05 | 0.82 |
| | ***QLD n*** | ***NT n*** | ***WA n*** | | | |
| *Proportion of time spent moving (between regions)* | 10 | 10 | 10 | Exposure (Toad v Control) | 0.74 | 0.69 |
| | | | | Worm length | 0.01 | 0.92 |
| | | | | Ambient temperature | 0.12 | 0.73 |
| | | | | Worm family | 0.09 | 0.95 |
| *Average movement speed (between regions)* | 8 | 10 | 8 | Exposure (Toad v Control) | 2.92 | 0.08 |
| | | | | Worm length | 0.23 | 0.64 |
| | | | | Ambient temperature | 3.24 | 0.09 |
| | | | | Worm family | 0.26 | 0.77 |
| *Effect of lungworm activity on infection attempts* | 56 (10) | 50 (10) | 60 (10) | Average lungworm speed | 1.13 | 0.29 |
| | | | | Toad region of origin | 1.67 | 0.43 |
| *Effect of toad region on successful infections* | 8 | 8 | 10 | ***Toad region of origin*** | ***14.47*** | ***< 0.01*** |
